# Increased Demographic Turnover and Isolation Precede Social Fragmentation in Asian Elephants

**DOI:** 10.64898/2026.09.24.754174

**Authors:** Anastasia E. Madsen, T.V. Pushpakumara, U. Sameera Weerathunga, Devaka K. Weerakoon, Shermin de Silva

## Abstract

Humans and other animals live in interconnected social communities that are essential for survival and well-being. In principle, social communities can persist despite changes in constituent members. However, gradually accumulating changes to membership could push systems past critical thresholds into alternate states. We investigated social network dynamics for 443 wild female Asian elephants over 12 years by tracking community membership and structure. Following accumulated demographic turnover, networks fractured into smaller and more isolated social communities. Connections were changed not only due to turnover of members but also rewired among remaining individuals. This provides the first empirical evidence of a social network state change in a long-lived mammal.

## Main Text

Social species rely on conspecifics for meeting basic survival requirements like accessing food (*1*), caring for young (*2, 3*), and managing stress (*4–6*). Many species exhibit fission-fusion social dynamics (*7, 8*), in which associations among individuals are influenced by resource availability (*9–11*), predation risk (*12, 13*), and demographic turnover (*14–18*). While social communities can in principle persist despite this flux, akin to the Ship of Theseus that maintains its structure and function despite the replacement of individual parts (*19*), unexpected or lasting changes in structure can signal that critical shifts have occurred in the underlying processes that drive individual social behavior (*20*). Such shifts may occur when populations decline below critical thresholds, analogous to demographic Allee effects (*21–23*). This raises a critical question: how resilient are social systems to demographic dynamics?

In complex systems, abrupt state changes can occur when surpassing tipping points, or critical thresholds, following the gradual accumulation of changes to underlying conditions (*24*). For example, increasing nutrient loads in freshwater lakes can trigger a definitive shift from clear to murky, algae-saturated waters (*25*). Social systems can similarly reach tipping points, characterized as sudden shifts in behavioral states (*26, 27*) or cultural norms (e.g. establishment of cultural variants; *1, 28*). However, shifts documented in non-human species to date are typically based on changes in *individual behavioral states* (*27*), not on changes in the state of *globalized social network properties*. Documenting the latter is challenging as it requires longitudinal, individual-based observations of large numbers of individuals, which are rare for many terrestrial vertebrates (*29, 30*). As a result, state changes in animal societies are not well documented or understood despite growing interest (*31–33*). Indeed, possible social state shifts have primarily been proposed in theoretical simulations, but have lacked empirical evidence (*34, 35*).

We present the first empirical example of a social state change following gradually accumulating demographic turnover in a long-lived mammal, wild Asian elephants (*Elephas maximus*). Typically, female elephants exhibit complex and multi-level societies that are mostly based on matrilineal kinship (*36–39*). However, many elephant populations are highly disturbed and experience heightened mortality due to poaching and conflict with people, impacting population dynamics and social behaviors (*40, 41*). While some populations can regain social structure and function after poaching events (*42*), others may lose sources of generational knowledge (*43–45*). Social changes have been linked to targeted losses of key individuals and punctuated disturbances (*45*), however, the potential social impact of gradual demographic decline has not been documented in any population (*23*). Here we apply network analyses to longitudinal social data to investigate the effects of demographic turnover on Asian elephant social structure.

### Social communities become smaller and more isolated following demographic turnover

The Asian elephant population passing through Udawalawe National Park, Sri Lanka has been monitored since 2006 and was estimated to number between 804-1160 individuals between 2007-2008. We evaluated the demographic change in the population by monitoring uniquely identified adult and subadult females between years from 2007-2018 (Figs. 1A,B). Annual deaths and permanent dispersals increased over time, quickly surpassing annual immigration after 2012 (Fig. 1C). This pattern was not merely due to changes in observation effort (Fig. S1). Accordingly, there was a reduction in the proportion of individuals that remained in the population from the previous year, and the number of individuals that were observed from the beginning of the study declined dramatically over the study period, with ∼70% of the original females lost by 2018 (Fig. S2) although elephants in this population are known to be capable of living 60 years or more (*46*). This loss reflects presumed and confirmed deaths as well as permanent dispersals.

**Fig. 1.**
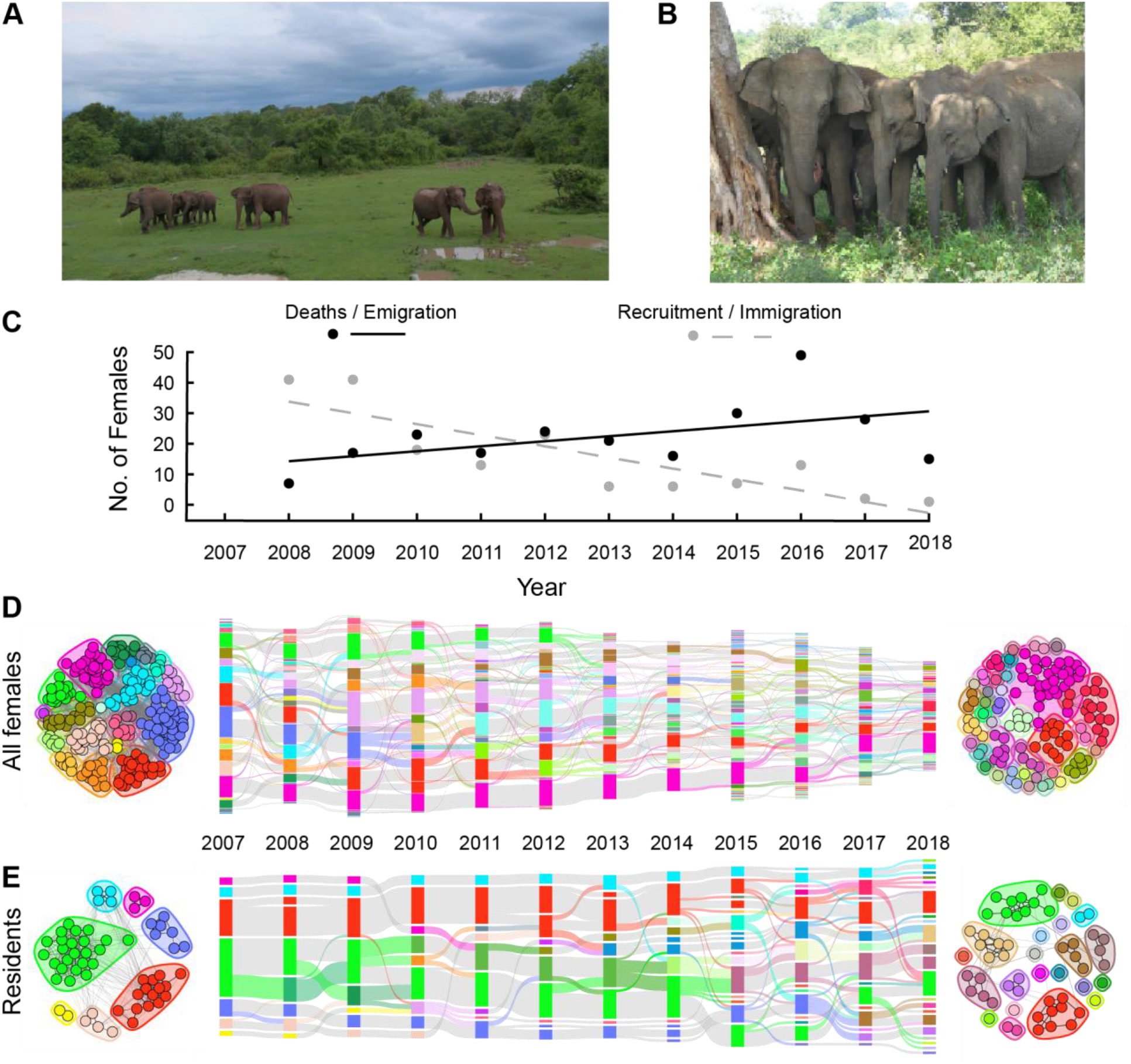
Social groups, demographic turnover and longitudinal dynamics in Asian elephants. Females and calves move and interact in small groups **(A)**, sharing resources such as water and shade (B). The number of deaths and permanent emigrations increases and the number of new immigrants decreases over time (C). The intersection of the regression lines occurs near 2012, after which emigration/deaths increasingly exceed immigration. Alluvial plots illustrate community dynamics for the full population (D), *N* = 443 individuals, 245 unique communities cumulatively, and residents (E), *N* = 59 individuals, 50 unique communities cumulatively. Vertical blocks represent distinct communities. Height is relative to the size of communities and the population in that year, relative to each graph. Horizontal bands represent the flow of individuals among communities over time: gray bands correspond to consistent membership and colored bands correspond to changing membership, colored by the source community. Some colors have been adjusted manually for visual clarity. Networks for 2007 (left) and 2018 (right) are colored by the corresponding community identity matching the alluvial plots for each set. Photographs in a-b by the Udawalawe Elephant Research Project.

The pattern of relationships (i.e. social structure; *47*) frequently results in dense clusters of connections among subsets of individuals, often called social communities (*48, 49*). The size and composition of social groupings (i.e., social organization; *50*) can vary over time due to changing individual preferences (*7, 51*) and demographic turnover (i.e. birth/immigration and death/emigration in the population; *14–18*). To investigate temporal changes to social structure and organization, we built annual social networks from spatiotemporal association data.

Associations were derived from co-occurrences of individuals observed moving together and exhibiting affiliative behavior (Figs. 1A,B; quantified with the Simple Ratio Index of association, SRI; *52, 53*). Adult females in this population change associates on a day-to-day basis but maintain long-term social affiliates over multiple years (*53*). We constructed networks that included adult and subadult females (“full female population”, Fig. 1D, *N* = 443) and networks for a subset of core “residents” that were present in all years of the study (Fig. 1E, *N* = 59). This allowed us to distinguish structural changes in networks independent of the population size (i.e. number of nodes). We identified social communities in each network using the Louvain community clustering algorithm, which detects multiscale organization from the pattern of connections across each annual network (*48, 54*), and tracked the membership in these communities between years (detailed in Methods; *55*). Clustering was run on networks for the full female population and residents separately. The detected communities were robust irrespective of the choice of clustering algorithm (Fig. S3-4). Mean community size decreased over time, accompanied by an increase in the mean number of communities (Fig. 2A,B). These changes to social organization were evident in the full female population immediately following the year in which emigration exceeded immigration (2012), but occurred after a lag of 1-3 years among the residents. Furthermore, the mean size and number of communities in the female population noticeably fluctuated prior to this shift, whereas they remained stable for the subset of residents (Fig. 2A,B).

**Fig. 2.**
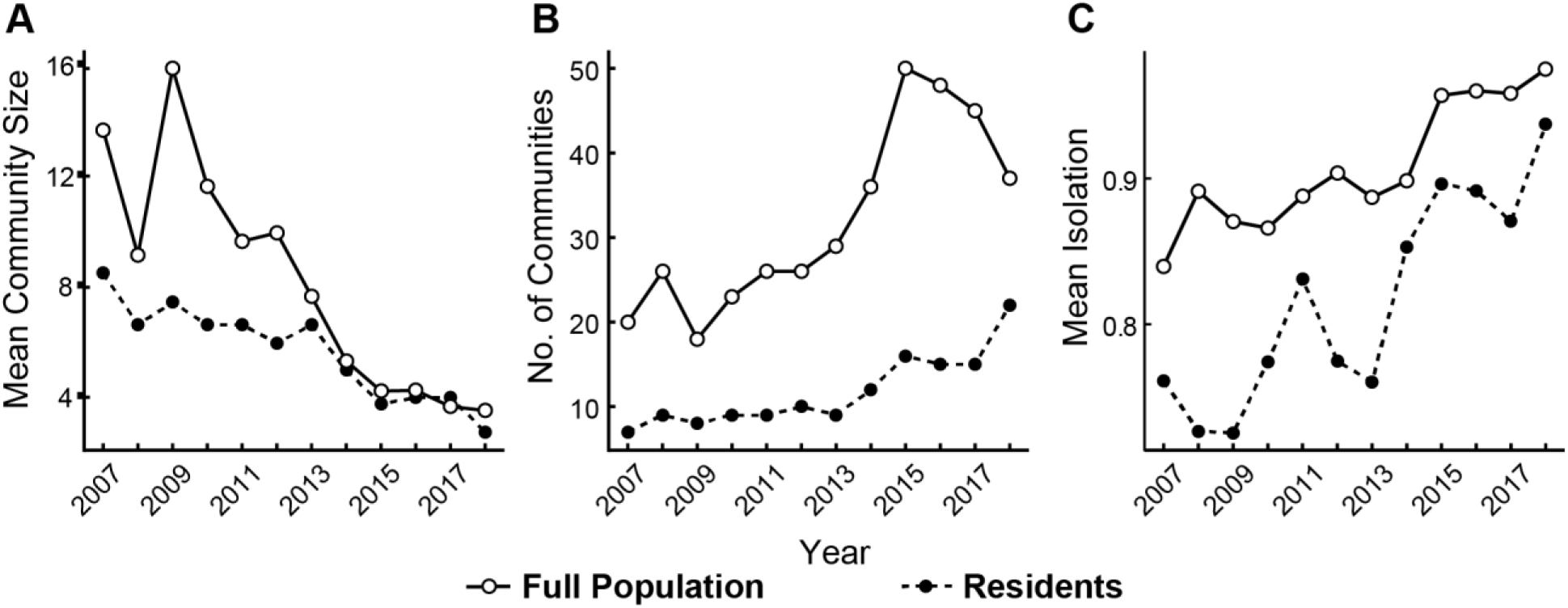
Changes to social community statistics over time. Over time, the mean size of these communities decreased **(A)**, the number of detected communities increased (B), and the structural isolation of communities increased (C). Structural isolation is defined as the proportion of connections that are contained within a community out of all connections of its members within and outside of the community, weighted by the Simple Ratio Index (SRI) values. Values of all statistics fluctuated prior to the state shift in 2012 for the full population (see also Fig. S5), but occurred after a shift in strict residents, except for isolation which spiked and briefly recovered prior to the shift (C).

Connections within and among social communities were used to evaluate community structure over time. Specifically, we assessed the degree of structural isolation of communities. Structural isolation was defined as the proportion of edges contained within a social community out of all connections of its members within and outside of the community, weighted by SRI values (Fig. 2C; *56, 57*). Greater structural isolation values indicate fewer connections between communities. There was a sudden increase and fluctuation in isolation among the residents in the years leading up to 2012, reflecting that communities started becoming more disconnected from others before further fragmentation (Fig. 2C). Subsequent trends were consistent for the full female population as well as residents, demonstrating that the observed fragmentation of communities in later years was not solely a function of population size, but reflects a restructuring of relationships among individuals. We calculated a range of statistics for each network that indicate stability and change in social networks, a majority of which showed a trend towards a less connected society over time (Fig. S5), with many correlating significantly with increasing turnover rates (Table 1, Fig. S6).

**Table 1.** Correlation of network statistics with turnover. Linear regression models show the correlation of each social network statistic with the proportion of the population that was lost to turnover (see also Fig. S6). Separate models were fit for the full population and residents. Significant two-tailed p-values are indicated in bold.

| Subset | Statistic | Estimate | SE | P |
| --- | --- | --- | --- | --- |
| Full Population | Edge Density | -0.0025 | 0.00079 | **0.012** |
| | Mean Normalized Strength | $8.68 \times 10^7$ | $3.85 \times 10^7$ | 0.051 |
| | Mean Node Betweenness | $1.56 \times 10^4$ | $8.68 \times 10^5$ | 0.11 |
|  | Mean Normalized Path Length | 0.023 | 0.0055 | **0.0021** |
|  | CV SRI | -0.025 | 0.0064 | **0.0035** |
|  | Global Clustering | 0.0034 | 0.0020 | 0.12 |
|  | Modularity | 0.013 | 0.0032 | **0.0031** |
|  | Mean Community Size | -0.48 | 0.16 | **0.016** |
|  | No. of Communities | 1.43 | 0.41 | **0.0064** |
|  | Mean Structural Isolation | 0.0050 | 0.0015 | **0.0077** |
| Residents | Edge Density | -0.011 | 0.0038 | **0.018** |
| | Mean Normalized Strength | $9.11 \times 10^6$ | $2.4 \times 10^6$ | **0.0046** |
|  | Mean Node Betweenness | 0.00059 | 0.00048 | 0.25 |
|  | Mean Normalized Path Length | 0.030 | 0.0056 | **0.00049** |
|  | CV SRI | -0.036 | 0.0072 | **0.00073** |
|  | Global Clustering | -0.0073 | 0.0035 | 0.065 |
|  | Modularity | 0.018 | 0.0055 | **0.011** |
|  | Mean Community Size | -0.20 | 0.062 | **0.012** |
|  | No. of Communities | 0.51 | 0.19 | **0.024** |
|  | Mean Structural Isolation | 0.0084 | 0.0034 | **0.037** |

### Community restructuring is driven by loss of nodes and indirect rewiring of weak ties

To identify the structural changes that accompanied community fracturing, we evaluated changes to network edges over time, first testing whether edge weight impacted the probability of edge rewiring between remaining individuals (i.e., conserved nodes) in consecutive years. For the full population, we excluded direct effects of node turnover by only considering edges between individuals that were conserved between each pair of consecutive years. Rewired edges were significantly weaker than retained edges, with a sharp drop in rewiring probability for edge weights greater than 0.5 (binomial generalized linear model, full population *N* = 335, MLE = - 8.0988, SE = 0.1596, *z* = -50.74, *p* << 0.001, Fig. 3A; residents, *N* = 59, MLE = -10.9975, SE = 0.4618, *z* = -23.812, *p* << 0.001, Fig. 3B). We also tracked the number of edges that were conserved, edges that were gained or lost due to turnover, and edges that were gained or lost due to rewiring, which we show between consecutive years (Fig. 3C,D) and from the beginning of the study (Fig. 3E,F). While these proportions were relatively constant between consecutive years (Fig. 3C,D), edges that changed due to turnover gradually increased over time for the full population (Fig. 3E,F), as expected from the overall population trend. Rewiring increasingly led to the loss of edges for both the full population and residents (Fig. 3E,F), indicating that remaining individuals were less likely to re-associate with previous companions over time (Fig. S7). The direct effects of turnover and indirect effects of increased rewiring leads to increasing instability of weak ties over time.

**Fig. 3.**
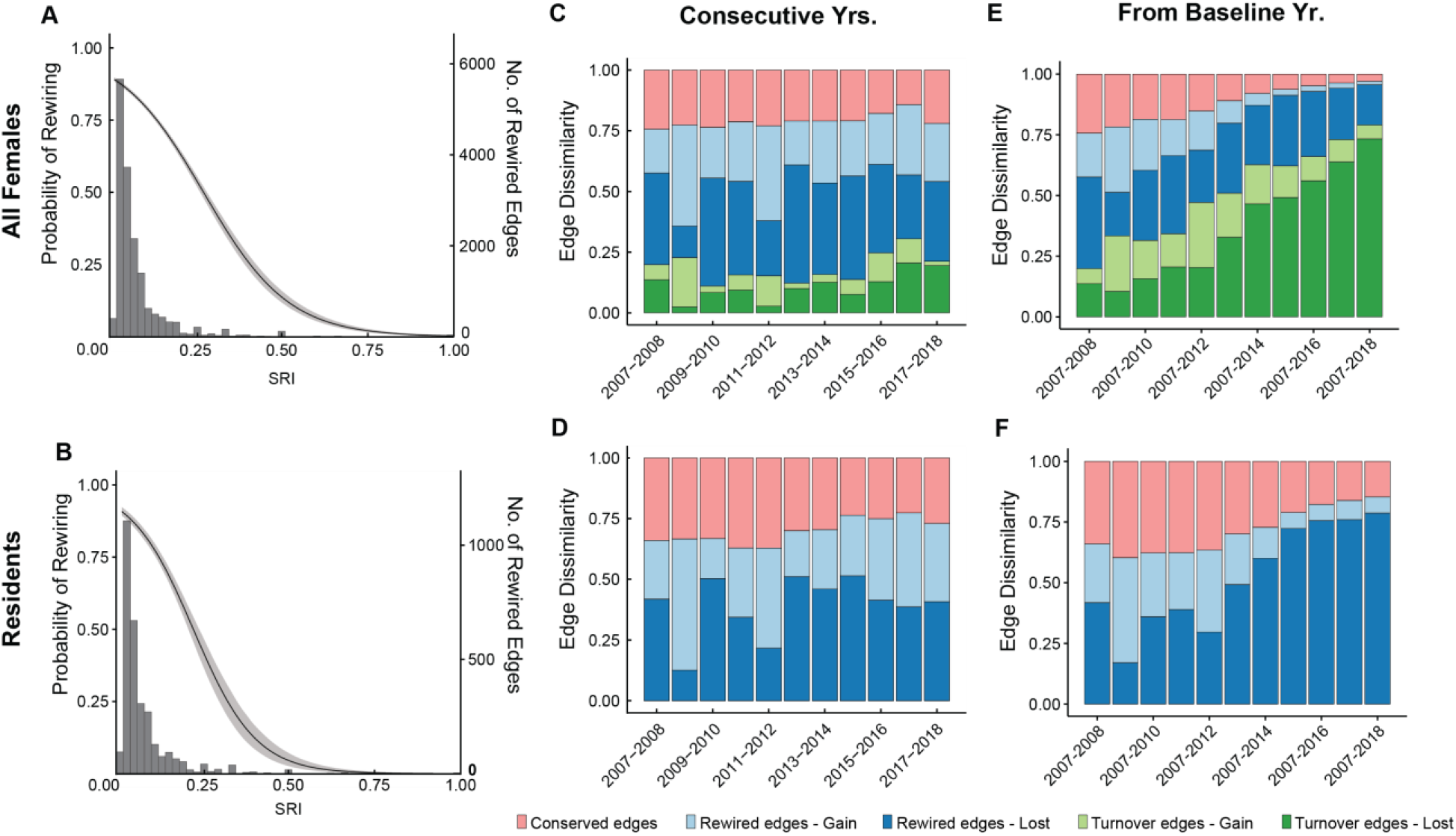
Structural changes due to direct and indirect effects of turnover. The probability of edge rewiring between years by edge weight value (Simple Ratio Index, SRI) is shown for all individuals in the full population that were present in pairs of consecutive years (A), *N* = 335 and the residents (B), *N* = 59. Curves show the output of a binomial regression with a logit-link function and gray bands represent bootstrapped confidence intervals (a, MLE = -8.0988, SE = 0.1596, *z* = -50.74, *p* << 0.001; b, MLE = -10.9975, SE = 0.4618, *z* = -23.812, *p* << 0.001). Histograms **(A**,**B)** show counts of rewired edges at a given SRI value. Edge dissimilarity **(C-F)** represents the proportion of edges in networks that were conserved, lost or gained due to rewiring (i.e., between conserved individuals), and lost or gained to turnover (i.e., individuals entering and leaving the population). Edge dissimilarity is shown between consecutive years **(C**,**D)** and with respect to the baseline year 2007 **(E**,**F)**, for the full population (C,**E**; *N* = 443) and residents (D,**F**; *N* = 59). Gains are shown in lighter shades and losses in darker shades.

### Multi-level social structure and organization are changed but not eliminated

Elephant societies have multi-level structure, with this population previously being shown to exhibit at least two structural levels (*36, 53*). We therefore tested whether the observed fragmentation reflected a complete collapse of multi-level structure (namely, loss of higher-level communities maintained by weak connections), or more subtle re-structuring through the re- distribution of edges. Structural changes were evaluated by sequentially removing edges below an edge weight (SRI) threshold at 0.01-unit increments and re-clustering at each step (*36, 53*). Discontinuities in the resulting curves are indicative of regions of structural change. We found that discontinuities occurred at SRI thresholds between 0.25-0.5 for the full female population in all years (Fig. 4A,B), but for residents these were in fact evident only after 2012 (Fig. 4D,E). Surprisingly, this indicates that although the original larger higher-level communities broke down, networks still exhibited multi-level structure (Figs. S8-S9). Furthermore, when we quantified the distributions of edge weights over time, the weakest edges (SRI<0.2) became less common over time, but intermediate values (SRI ∼0.3) became proportionally more common even as networks became more sparse (Fig. 4 C,F).

**Fig. 4.**
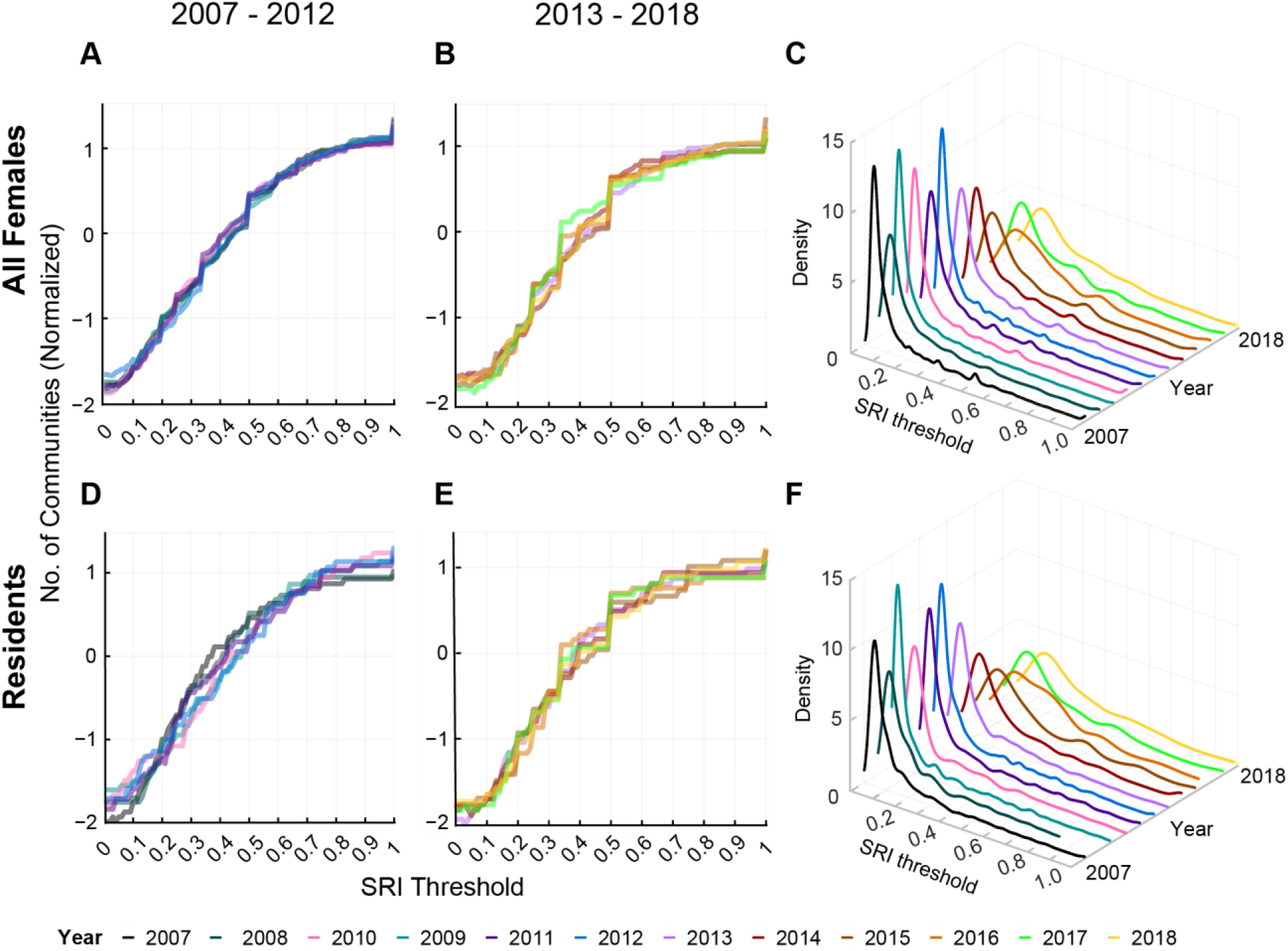
Higher-level social structure and organization are modified over time. The number of community clusters are shown over an increasing threshold of edges being removed from the weakest to strongest edge weight value (Simple Ratio Index, SRI; panels **A-B**,**D-E**). The size of networks differs between years, so the number of communities is normalized here to compare the placement and magnitude of structural changes in response to edges being filtered. The data is separated into panels by year (A,D: 2007-2012, B,E: 2013-2018) and network subsets (top row: full population, bottom row: residents). The density of SRI values for the full population (C) and residents (F) in each year shows that very weak connections gradually become less common over time and intermediate values become more prominent.

## Discussion

Demographic turnover may induce social state shifts by changing multiple aspects of network structure and organization, including population size, group composition, interaction rules, and mechanisms of integration (*58*). We find that gradual accumulation of losses due to turnover preceded sudden change in the social structure and organization among females in a wild population of Asian elephants. We propose that when the net balance of immigration and emigration became negative, the population passed a critical threshold (social tipping point), shifting to a new social regime characterized by smaller and less connected social communities (Figs. 1-2, Figs. S5-S6, S9, Table 1). This shift was preceded by increased isolation and fluctuations in network statistics (Fig. 2, Fig. S5), which is indicative of the system entering an unstable transition state (e.g., “flickering,” *59–61*). Furthermore, the shift was evident shortly after the tipping point at the scale of the full female population but was more subtle among more residential individuals, with a lag of 1 to 3 years before effects became pronounced (Figs. 1-2, Figs. S5-S6, S9, Table 1). This suggests that one or more underlying variables are dependent on prior states of the network. Increased variability, lagged responses, and shifts between alternate states also suggest the possibility of hysteresis in the system, where there are multiple possible states for a given set of values in underlying variables (*24, 59*).

Subsequently, larger higher-level communities fragmented into smaller components primarily due to the loss of very weak connections over time (Figs. 3-4, Figs. S8-S9), not only due to turnover but even among individuals that were present throughout. This demonstrates that the impacts of demographic decline can be both direct (due to gain/loss of nodes; *14, 62*) and indirect (gain/loss of edges among conserved nodes; *15, 17, 63–65*). This could be because some certain nodes mediate connections among multiple individuals. The population nevertheless retained multi-level structure and organization through increased associations of intermediate strength, even as the overall edge density degraded over time (Fig. 4, Fig. S9). This reflects that females are consolidating their relationships and selectively associating with only their strongest companions. To our knowledge, this is the first empirical example of a persistent state change in the overall social network structure of a long-lived non-human species.

Social connectivity influences many processes in societies with fission-fusion dynamics (*8, 51*). Cross-community connections can determine the trade-offs between affiliative and competitive behaviors (*66, 67*), while within-group connections can reinforce cooperation and cohesion (*68–71*). African savannah elephants show fission-fusion behaviors between seasons but maintain stable, long-term hierarchical relationship among matrilines that mediate competition and resource access (*37, 38, 42*). Asian elephants in southern India and Sri Lanka show more fluidity than African savannah elephants, but changes in association are also likely driven by seasonal resource availability (*53, 72, 73*). Further study is required to disentangle the demographic and ecological mechanisms driving these patterns. One possible explanation is that network fragmentation is due to the loss of key individuals, such as older adult females (*74, 75*). Alternately, changes in the local ecosystem, such as depletion of forage, may have caused individuals to die or disperse as well as forage more independently. These mechanisms are not mutually exclusive (*76–78*).

In a rapidly changing world, we need a clearer understanding of why animal societies undergo structural shifts and how to detect them. Weak connections can act as structural redundancies that buffer any type of network against some degree of perturbation (*79, 80*). Reduced spatial density can trigger nonlinear losses of social connections (*81*), weakening resilience (*79, 80*) and hastening demographic collapse (*21–23*) through feedbacks between social and demographic processes (*82–85*). Moreover, social relationships take time to form and develop (*32, 86, 87*), an important consideration for long-lived species with slower life histories (*88, 89*). As connectivity erodes and groups shrink, societies may experience reduced foraging efficiency (*90*), greater aggression(*91*), weakened herd immunity (*92*), and declines in overall fitness (*93*), alongside disruptions to cultural transmission (*94, 95*).

Disturbances or the loss of important relationships can disrupt cohesion (*75, 84*), information flow (*96*), and cultural norms (*97, 98*), with profound consequences for the well-being of individuals and social communities (*32, 99–102*) weakening their ability to recover from future disturbances. The loss of socially important individuals can further accelerate changes in norms (*97, 98*), intragroup interactions (*75*), and community structure (*14*). Although demographic factors are central to predicting social tipping points, these shifts are difficult to forecast and are especially underappreciated in slow-breeding, long-lived species whose longevity can mask gradual decline (*23, 103–105*).

The direction, magnitude, and timing of structural changes in networks may point to which interaction rules are changing (*33, 58, 106, 107*), allowing targeted interventions to encourage population recovery (*18, 108*). Identifying network statistics that could reliably act as early indicators of social state change could be of vital importance for species of conservation concern (*108*) but ideally requires many more studies across diverse taxa to quantify the normal range of variability within systems (*30, 76*). Such monitoring may be extremely difficult to achieve on practical timescales (*109*), especially when anthropogenic stressors are now ubiquitous (*110, 111*). Our observations highlight the need for a precautionary approach to conserving social species by recognizing that declining numbers may have delayed and unanticipated social consequences.

## Acknowledgments

The authors thank James Nieh, Jeff Lucas, and Sergey Kryazhimskiy for helpful comments on drafts of the manuscript and Colton Fruhling for assistance with 3D panels in Figure 4. We thank the Department of Wildlife Conservation Sri Lanka for granting permission to conduct this work (permits WL/3/2/1/6, WL/3/2/4/12).

## Funding

US Fish and Wildlife Asian Elephant Conservation Grant F11AP00258 (SdS) US Fish and Wildlife Asian Elephant Conservation Grant F14AP00256 (SdS)

US Fish and Wildlife Asian Elephant Conservation Grant F17AP00308 (SdS) National Science Foundation DBI-2305860 (AEM) https://credit.niso.org/ University of Pennsylvania

University of California San Diego Trunks & Leaves Inc.

## Author contributions

Conceptualization: AEM, SdS

Data Curation: AEM, SdS

Formal analysis: AEM

Funding acquisition: AEM, SdS

Investigation: AEM, SdS, TVP, USW, DKW

Methodology: AEM, SdS

Project administration: SdS, DKW

Supervision: SdS, DKW

Visualization: AEM, SdS

Writing – original draft: AEM

Writing – review & editing: AEM, SdS

## Competing interests

Authors declare that they have no competing interests.

## Data, code, and materials availability

The datasets analyzed during the current study and associated R coding scripts will be made publicly available in a Data Dryad repository (*112*).

## Notes

### Competing Interest Statement

The authors have declared no competing interest.

