## Supplemental materials for "Increased Demographic Turnover and Isolation Precede Social Fragmentation in Asian Elephants"

**Supplementary Materials**

**Authors:** Anastasia E. Madsen<sup>1\*</sup>, T.V. Pushpakumara<sup>2</sup>, U. Sameera Weerathunga<sup>2</sup>, Devaka K. Weerakoon<sup>3</sup>, Shermin de Silva<sup>1,2\*</sup>

**Affiliations:**

<sup>1</sup>Department of Ecology, Behavior and Evolution, University of California; San Diego, 92093 USA

<sup>2</sup>EFFECT; 215 A 3/7 Park Road, Colombo, 00500, Sri Lanka

<sup>3</sup>University of Colombo; Colombo, 00300, Sri Lanka

### Materials and Methods

#### Study System

Udawalawe National Park, southern Sri Lanka is a 308 km<sup>2</sup> protected area consisting of grassland, mixed evergreen and deciduous forest, scrubland, riverine forest and wetlands, including the Udawalawe reservoir (1). The park was previously estimated to contain a population of ~1000 elephants, with seasonal fluctuations as well as permanent immigration and emigration (1, 2). Herds of female elephants were observed from 2007-2018 between 0600 and 1830 throughout the year with 1665 total observation days and an average of 12.15 observation days per month (SD = 4.99, Fig. S1). To record observations of elephants, observers drove along road networks throughout the park. When a herd of elephants was spotted, a GPS point was taken from the road. All individuals within 500 meters of each other that were moving cohesively together, interacting affiliatively, and sharing resources (Fig. 1A,B) were considered to be associating in the same group (1). Individuals were identified using unique physical features from a catalogue of reference photographs (1). We focus our analyses on adult females because they were reliably identified throughout the study; subadult females that were likewise reliably identified during the study period were also included in our analyses (total $N = 443$).

#### Demographic Turnover

Population size trends were tracked over time to assess the number of individuals entering the population for the first time and individuals leaving the population permanently due to deaths or emigration and the rate of turnover (Fig. 1C & Fig. S2). The rate of turnover in the population was evaluated by tracking the number of elephants that left, remained, and entered the population over time. This included consecutive-year changes, which is the interannual turnover rate, and changes from the first year of the study (2007 baseline), which shows the extent of individuals leaving the population (Fig. S2). Specifically, we calculated the relative turnover rate as $(E+L) \times 100 / (N_i + N_{i+1})$, where $E$ was the number of new individuals entering the population, $L$ was the number of individuals leaving the population, $N_i$ was the number of individuals in year $i$ and $N_{i+1}$ was the number of individuals in year $i+1$ (mean = 33.78%, SD = 5.77%). (3) We also evaluated turnover from the first year of the study, calculated as $(E+L) \times 100 / (N_1 + N_{i+1})$ to show accumulating differences in turnover (mean = 54.78%, SD = 13.13%).

#### Social Networks

Networks were built annually using the Simple Ratio Index of association, which represents the likelihood that any pair of individuals were seen together out of the total sightings for both individuals together and apart (SRI) (4). Asian elephants range widely and are capable of communication over long distances, so we calculated SRI using a sampling period of one day to account for the possibility that subsets of groups sighted separately were maintaining cohesion at a spatial level not detected in observers' field of view (1). Because individuals join and leave the population over time due to different demographic processes, we calculated networks for the entire population ($N = 443$ total individuals) and for a subset of the population that was present in every year of the study, which we will refer to as "residents" ($N = 59$ individuals), though we note that these individuals can also occasionally or regularly disperse outside the park. Comparing changes in network statistics with changing population size to those from networks of a constant size allows us to distinguish whether trends are dependent on population size alone.

### Social Community Detection

#### *Clustering Algorithm Comparison*

We chose the Louvain clustering algorithm to identify community structures in networks to maintain consistency with other published studies on Asian elephant social behavior(5, 6). This algorithm uses a hierarchical clustering method that optimizes modularity, which is the density of edges inside communities to edges between communities, while accounting for edge weights.(7) To ensure that trends observed in the data were not due to analytical artifacts, we compared the performance of the Louvain algorithm and three additional clustering algorithms available in the R package *asnipe*: Louvain (7), Girvan-Newman (8), Infomap (9), and Walktrap (10). For algorithms that are stochastic (i.e., dependent on the starting point), we ran the clustering for each method over 100 iterations for each annual network for the full female population and residents. We assessed the agreement of clustering algorithms across iterations by recording the number of communities, mean size of communities, modularity, and structural isolation of communities (aka, structural cohesion; see below for full explanation). All community statistics maintained a similar temporal trend regardless of the algorithm used (Fig. S3).

We next evaluated the similarity in membership assignment to the Louvain algorithm using the Rand Index, which ranges from 0 (no agreement in community assignments) to 1 (complete agreement in community assignments;11). Membership assignments were consistently similar between the Louvain and Infomap algorithms, and between the Louvain and Walktrap algorithms (Fig. S4). The output from the Girvan-Newman algorithm appeared overall to have less agreement with Louvain compared to other algorithms (Fig. S4). The Girvan-Newman method also does not consider edge weights in its calculations, but it is important to consider weights in the topology of a network to accurately detect community structures in real networks (12). As the trend of interest was the same regardless of algorithm and that we had high agreement among multiple independent algorithms, we continued with the Louvain algorithm for subsequent analyses.

#### *Long-Term Community Membership Dynamics*

To assess and visualize long-term community membership and dynamics, we applied a dynamic community detection algorithm to the Louvain clustering assignments (*Majortrack* (13)). This algorithm assesses the identity of communities between consecutive time points using a mutual majority of membership; i.e., to be identified as the same community between years, a community in year $i$ must contain a majority of the same members as a community in year $i+1$ and the community in year $i+1$ must contain a majority of the same members as the community in year $i$ (13). We configured the algorithm to look across all annual networks before determining community identity. This contrasts with the Louvain clustering, which only considers relationships within a given year. Using the results of this algorithm, we discerned whether a community persisted across time points (i.e., maintained its identity), broke apart into separate communities, fused into a community with a new identity, or disappeared when all members left the population (Fig. 1B-C). Importantly, community identity is used here to characterize the similarity in community assignments between consecutive time points, but these communities are not a priori assumed to be static over time, and network community identity derived from network topology does not correspond to spatiotemporally cohesive aggregations observed in the field. Assigning identity allows us to track the flow of members between community clusters over time as a measure of upper-level network structure and organization beyond those observed day-to-day. Here, we use this algorithm purely to visualize these changes to membership whereas the Louvain results were used for all quantitative analyses. As such, some community colors in the alluvial plot were adjusted manually to reflect distinct within-year membership assignments from the Louvain clustering algorithm (Fig. 1B,C).

#### Analysis of Social Structure and Organization

We evaluated structural and organizational components in each year for social communities, including 1) mean community size, 2) the number of communities, and 3) mean community isolation (Fig. 2). Isolation refers to structural connectivity between communities, defined as the proportion of edges between community members out of all connections of its members, weighted by SRI (aka, structural cohesion (14)). This represents the strength of boundaries between communities, where high values correspond to a community that is less connected (i.e., more isolated) from others and low values correspond to a community that is more broadly connected. Additionally, we evaluated additional global network statistics over time (Fig. S5) and the correlation of all community and global statistics with relative turnover rate (Table 1, Fig. S6). We selected network statistics that signal changes in social stability and network position (edge density, normalized degree strength (15, 16); node betweenness centrality (17)), connectivity (path length), heterogeneity among connections (CV SRI (4, 18)), and clustering of connections in the network (global clustering (19, 20), modularity (21)). Separate linear regression models were run for each statistic, where relative turnover rate (see above) was the predictor variable and the network statistic was the response (Table 1, Fig. S6). Regressions were implemented using ordinary least squares estimation in the *stats* R package (22). Finally, we assessed the stability of pairwise associations using lagged association rates across years (LAR), which is the probability of two individuals re-associating over increasing lengths of time (4, 23, 24). For both the full population and the residents, we calculated probabilities by year to represent annual networks and used a jackknifing approach to calculate standard error values (Fig. S7).

We tested whether the strength of connections (i.e., edge weight, SRI) impacted their probability of rewiring between consecutive years using a binomial generalized linear mixed effects model with a logit-link function (*lme4* R package; 25) and significance was evaluated with two-sided p-values. We initially included an interaction effect between year and SRI, but this model was overfitted and we removed the interaction effect in the final model, instead including the year as a random effect. This model was applied separately to the full female population and resident networks (Fig. 3A,B). The full population networks included only edges between individuals that were present in both networks being compared, isolating the edges that were rewired as opposed to those that were lost or created due to individuals joining and leaving the population, or edges that were maintained between years (i.e., between $N = 335$ individuals). We also assessed structural changes between years by calculating edge dissimilarity, which was the proportion of edges that were maintained, lost or gained to rewiring among conserved individuals, and lost or gained to turnover between consecutive years (Fig. 3C,D) and from 2007 (Fig. 3E,F).

To evaluate changes to multi-level social structure stemming from the structure of edges in the network, we removed edges for each annual network in sequence by thresholding the SRI values from 0 to 1 in 0.01 increments (1, 26). In each step, we removed all edges below the threshold, re-ran the clustering algorithm, and recorded the number and the mean size of communities (Fig. 4A-B,D-E, Fig. S8). To evaluate changes to the distribution of SRI values over time, we calculated kernel density estimates (*stats* R package; 22) over all edges for each annual network for the full population (Fig. 4C) and the residents (Fig. 4F).

We additionally compared changes in community membership for upper- and lower-levels of organization from the Louvain clustering community assignments (Fig. S9). The Louvain algorithm uses a hierarchical approach to identify community clusters in which the output of each iteration represents a different level of hierarchical clustering (7). For each subset of the population, the full female population and residents, the algorithm identified two levels of organization in all years. We show a visualization of these two levels for the residents for illustrative purposes (Fig. S9).

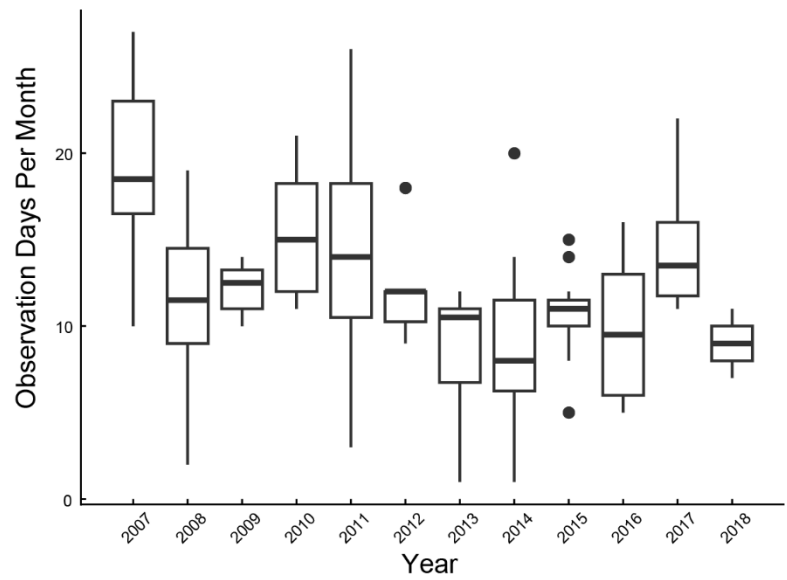

**Fig. S1. Number of sample days per month in each year.**

Boxes bracket the first and third quartiles, with the mean marked as a bold line. The ends of the whiskers extend to the largest value, except values outside the inter-quartile range, which are represented as outlier points. Mean = 12.15, SD = 4.99, $N = 1665$ total observation days.

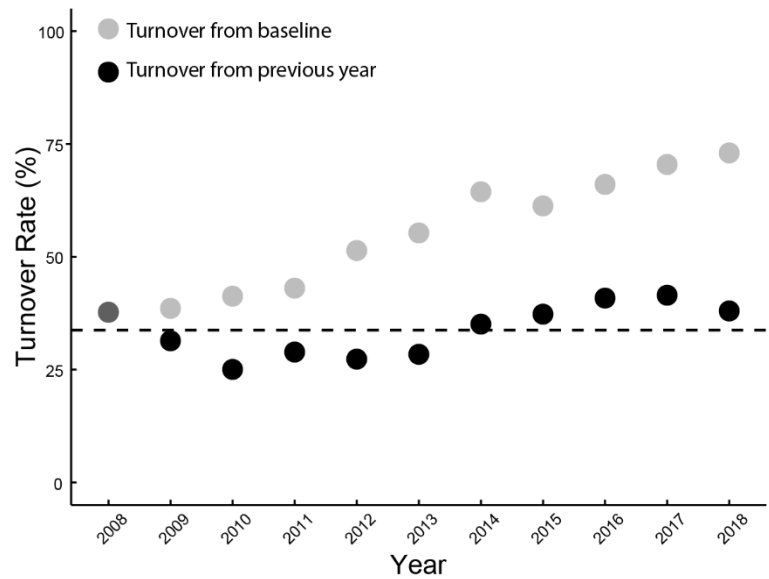

**Fig. S2. Female turnover in networks.**

Turnover rate is shown as the aggregate changes to network size due to immigration, temporary emigration, permanent emigration, or death. Turnover between consecutive years are shown as black points and proportions from the 2007 baseline are shown in gray (the first point is the same for both). The dashed line shows the mean turnover rate for consecutive years (mean = 33.78%; $N = 443$ total females).

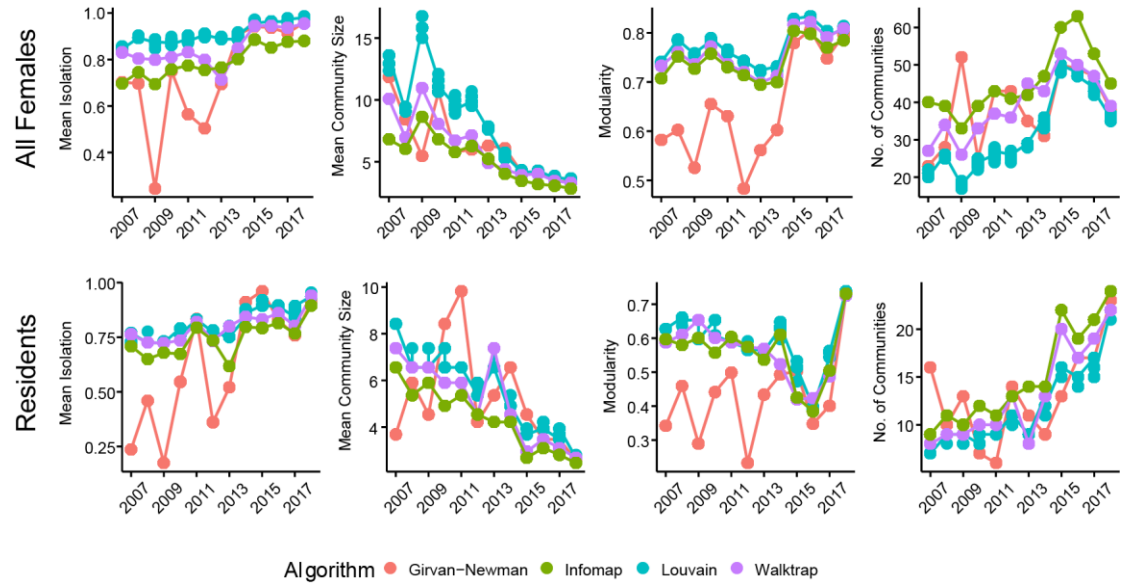

**Fig. S3. Comparison of statistics between community detection algorithms.**

Network statistics that rely on community detection methods showed similar temporal trends over time for all clustering methods. clustering was run over 100 iterations at different starting points.

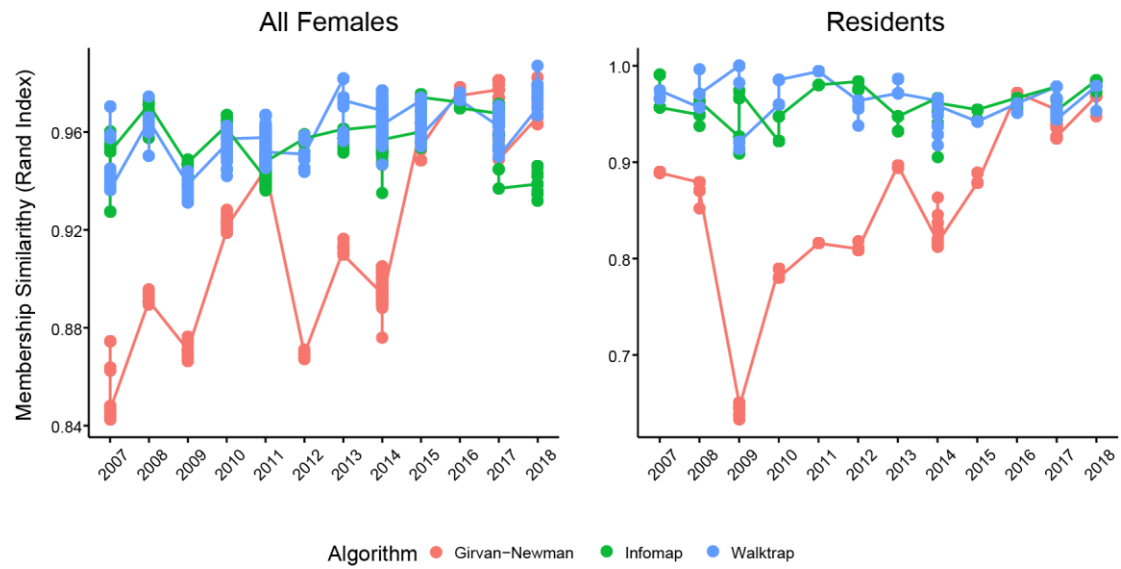

**Fig. S4. Comparison of membership assignments to the Louvain algorithm.**

Membership similarity (Rand Index) was calculated between the Louvain algorithm and Girvan-Newman, Infomap, and Walktrap. Similarities were calculated between algorithms for each clustering output over 100 iterations of different starting points.

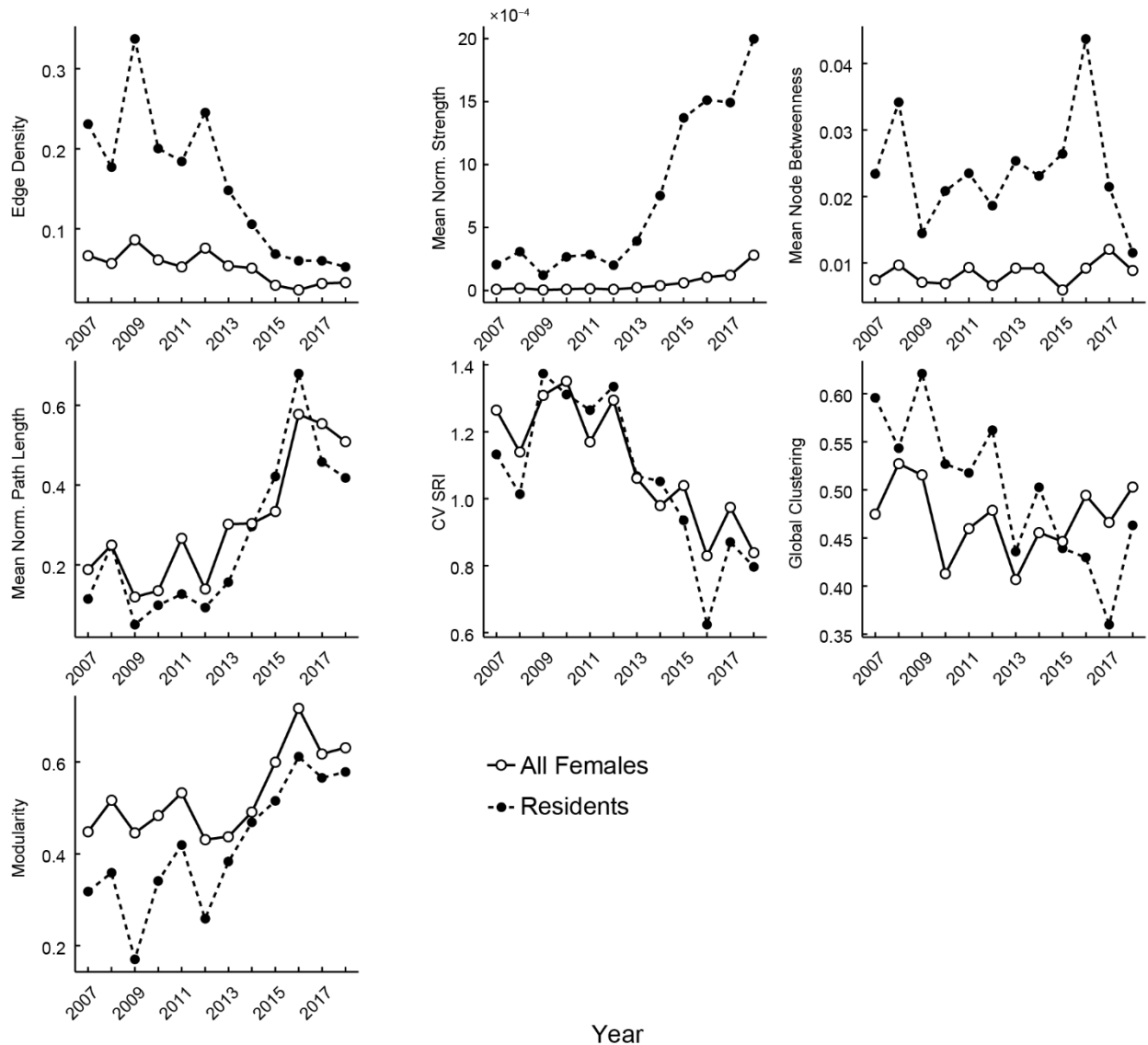

**Fig. S5. Network statistics over time.**

Trends in statistics are shown for the full female population (solid line, open circles) and residents (dashed line, closed circles). Both sets show similarly abrupt change after 2012, aside from mean normalized strength and mean node betweenness, each of which changed more gradually for residents.

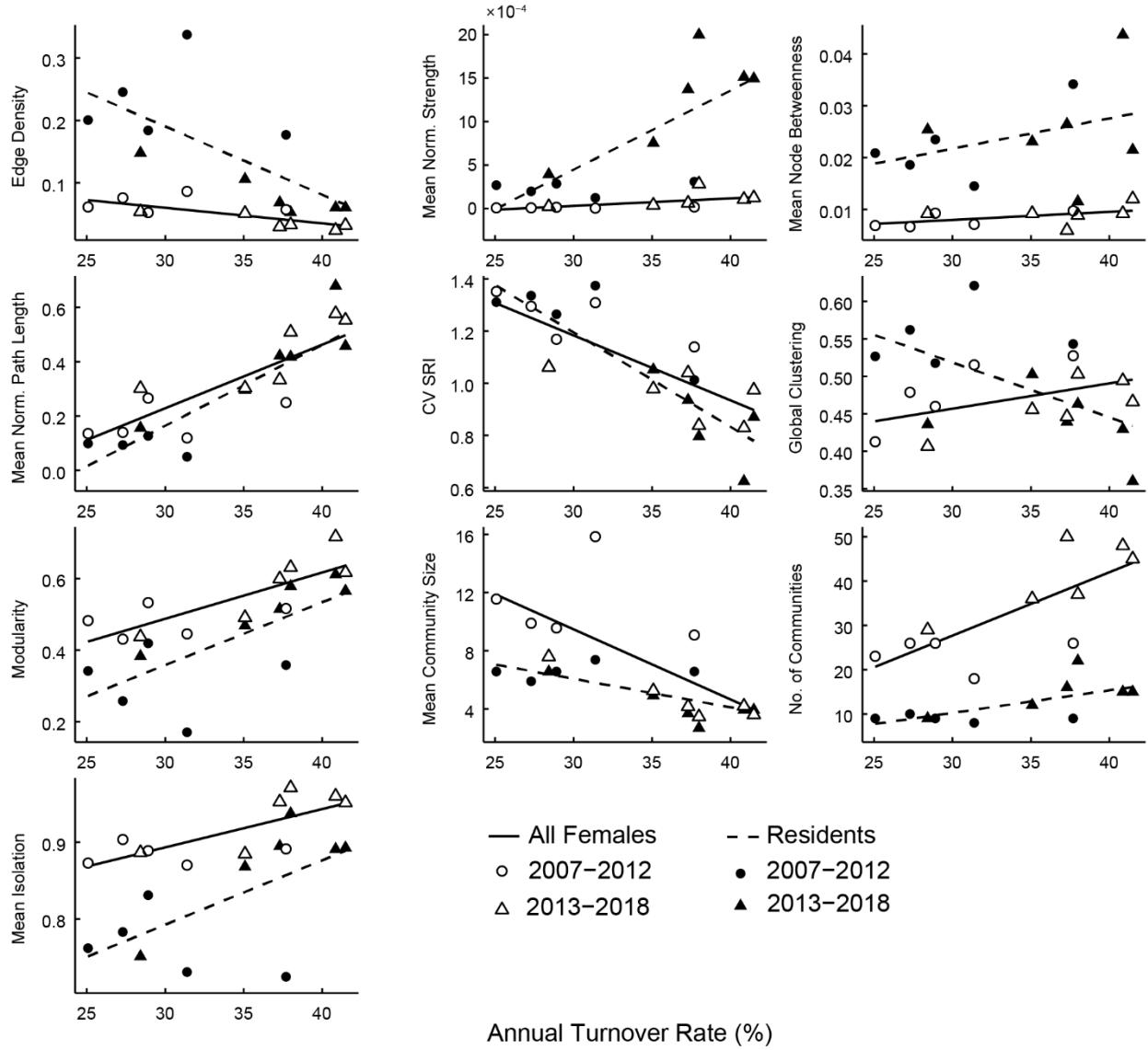

**Fig. S6. Changes in network statistics vs. turnover.**

Social structure and organization change with increasing turnover. Residents and the full population show similar trends for most measures, aside from global clustering ($N = 12$ points for each regression line).

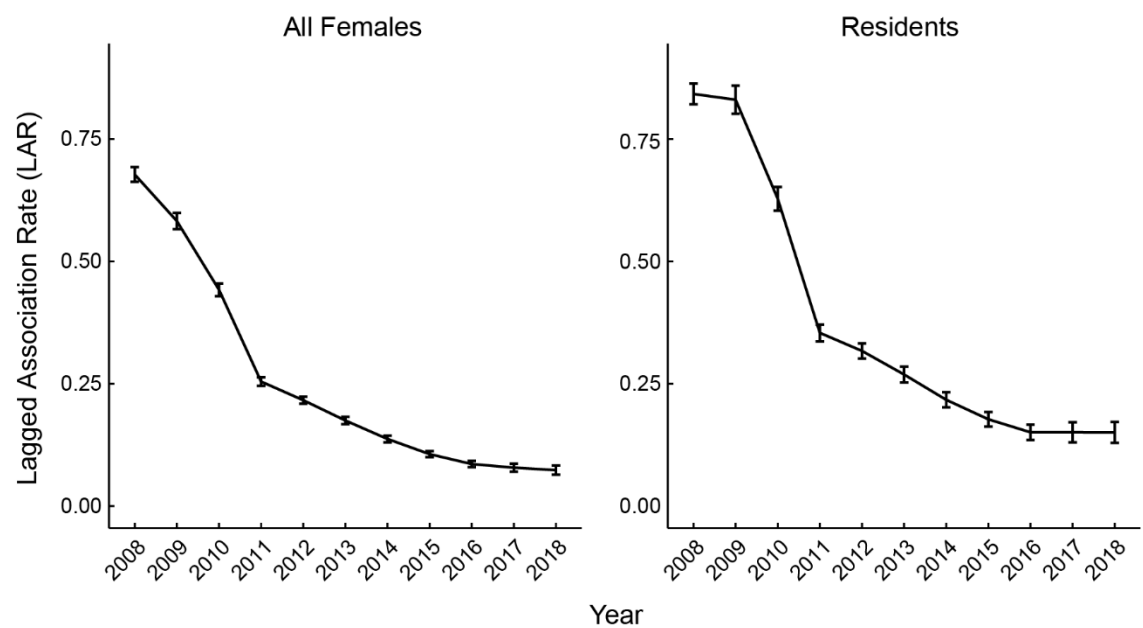

**Fig. S7. Lagged association rates over time (LAR).**

Associations among pairs of individuals became less correlated over time for the full population ($N = 443$) and residents ($N = 59$). Error bars show the standard error, calculated by permuting the raw data stream.

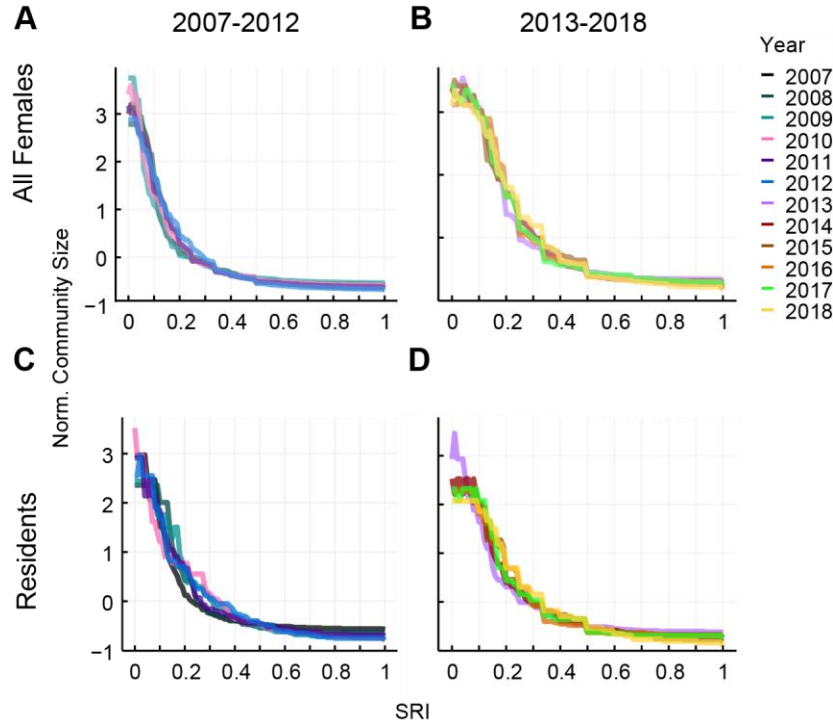

**Fig. S8. Higher-order social structure and organization breaks down over time.**

The data is separated into panels by year (left: 2007-2012, right: 2013-2018) and network subsets (top: full population, bottom: residents). Curves show the size of community clusters over when edges of a given weight (Simple Ratio Index, SRI) are sequentially removed. The sizes of networks differ between years and between each subset of the population, so community size is normalized to compare the placement and magnitude of structural changes in response to edges being filtered. As networks become sparser (b,d) the starting size communities declines. This is most evident among residents, where normalized community size starts at above 3 in 2013 but drops sharply in subsequent years.

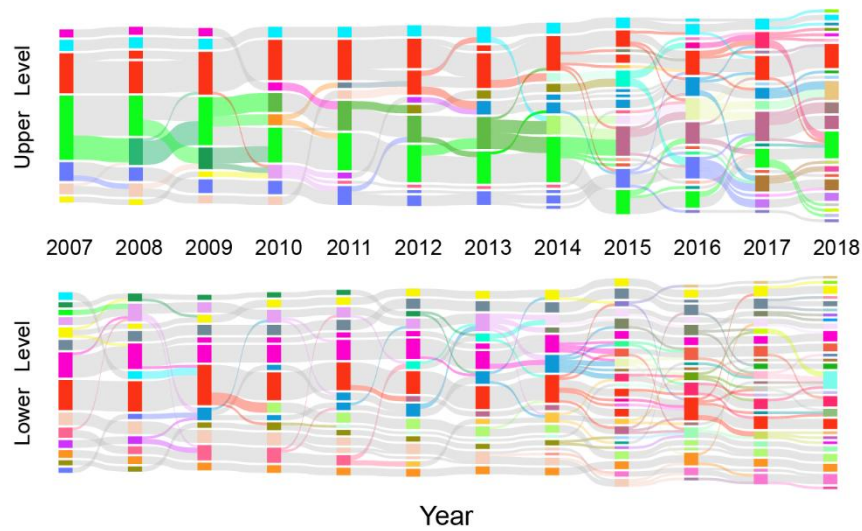

**Fig. S9. Alluvial plots showing Louvain clustering at two hierarchical levels.**

The upper-level plot is the same as in Fig. 1c in the Main Text, provided here for ease of comparison. Colors between plots are distinct. Vertical blocks represent distinct communities. Height is relative to the size of communities and the population in that year, relative to each graph. Horizontal bands represent the flow of membership over time: gray bands correspond to consistent membership and colored bands correspond to changing membership, colored by the source community.
